# First report of an insilico study of Luciferase in *Mycobacterium sp. EPa45*

**DOI:** 10.1101/2020.10.30.362764

**Authors:** Rishav Dasgupta, Arpit Kumar Pradhan, Shyamasree Ghosh

**Author notes:** Equal contribution (Rishav Dasgupta, Arpit Kumar Pradhan).

## Abstract

Mycobacterium are a genus of Actinobacteria known to be responsible for several deadly diseases in both humans and animals, including tuberculosis. Luciferase is the primary protein in *Mycobacteria* that plays a role in bioluminescence. It also plays a role in some bacteria of being a source of energy transference, such as in the case of lumazine proteins. Although studies have been conducted in different luciferase in bacteria, there has been hardly any structural studies on luciferase expressed in *Mycobacterium sp. EPa45*. Therefore, in this paper we have studied luciferase expressed in *Mycobacterium sp. EPa45* by insilico analysis of its structure from its protein sequence. We report the observed differences within luciferase reported from other strains of mycobacterium and pathogenic and non-pathogenic forms of bacteria in terms of their (i) physiochemical characteristics, (ii) protein structure, (iii) multiple sequence alignment and (iv) phylogenetic relationships. We report for the first time the relation of this specific strain of Luciferase in mycobacterium and bacterium at large.

**Highlights:**

- *Mycobacterium sp. EPa45* shows similar characteristics to pathogenic mycobacterium
- Analysis of Luciferase sequence and protein qualities provides insight to pathogenicity
- The deadly nature of infectious mycobacterium, especially with luciferase sequences similar to Mycobacterium *sp. EPa45*, is analyzed

## 1. Introduction

*Mycobacterium* sp are a member of the Actinobacteria genus, which share both bacterial and fungal qualities^1^. These are known to include some very deadly pathogens including *Mycobacterium rhodesiae* (*M rhodesiae*) which is a scotochromogenic pathogen isolated from certain Rhodesian patients with pulmonary diseases causing a poor prognosis, leading to death, and pathogenic form of *Mycobacterium tuberculosis* (*M tuberculosis*), known to cause tuberculosis^2 3^. Certain other deadly species of Mycobacteria includes *Mycobacterium leprae* (*M leprae*) which is known to cause leprosy and *Mycobacterium ulcerans* (*M ulcerans*) which is known to cause Buruli ulcers^4^. According to the World Health Organization (WHO), these diseases are still quite widespread, contrary to certain perceived beliefs. Tuberculosis remains the world’s most infectious virus, claiming 1.5 million annually,^5^ while 14 countries, mostly from the developing world, bear the brunt of the leprosy epidemic. The Buruli ulcer disease is also quite deadly, more than half of the patients diagnosed with it are under 15^5^.

*Mycobacterium sp. EPa45* is reported as aphenanthrene degrader, isolated from a phenanthrene-degrading consortium with a complete genome of 6.2-Mb single circular chromosome reported to contain a phenanthrene degradation pathway^6^. Bioluminescence is mediated by enzymes such as firefly luciferase and bacterial luciferase. Luciferase enzyme in Mycobacteria species is responsible for its bioluminescence. The scotochromogenic characterization *M rhodesiae* indicates that it develops pigmentation in darkness. Although luciferase is known for its role in biolumuniscence, studies on *Mycobacteria* has been mainly focused on assessments of immunity to mycobacterial infection of which some use the bioluminescent qualities of *M rhodesiae* to study parameters of immunity^3^. The bacterial luciferase is reported to be structural dimer, around 80 kDa, with an α- and β-subunit. It functions by catalyzing the long-chain fatty acids (>7 carbons) oxidation and riboflavin reduction, leading to emission of blue-green light (λ = 490 nm).

The reaction is as follows

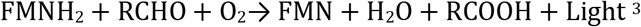

Bioluminescence, as produced by luciferase-catalyzed reactions, is widely used in the non-invasive localization of luminescent reporters in mycobacteria as reported from *Mycobacterium tuberculosis*. (*M tuberculosis*)^7^.

Although bioluminescence and luciferase has been studied in different groups of bacteria including those in the *Mycobacteria* species, there has been no structural evidence on the luciferase of *Mycobacterium sp. EPa45*. In this particular study, we have focused on understanding the structure of luciferase from *Mycobacterium sp. EPa45*, and make a structural comparison with luciferase found in pathogenic and non-pathogenic luminescent bacteria. For the pathogenic strain we have taken luciferase from *M tuberculosis*, and for the non-pathogenic strain we have taken luciferase sequence from *Escherichia coli*.

In this paper we study (i) the structure (ii) the physico-chemical and biological properties (ii) comparison with other similar proteins and function of luciferase *Mycobacterium sp. EPa45* from its sequence by applying *insilico* approaches.

## 2. Materials and methods

### 2.1 Sequences

All of the peptide sequences in this paper, including those of the related proteins to our strain of luciferase in question, were accessed via NCBI in the format of FASTA sequences. These sequences can be found in Table 1.

**Table 1:**
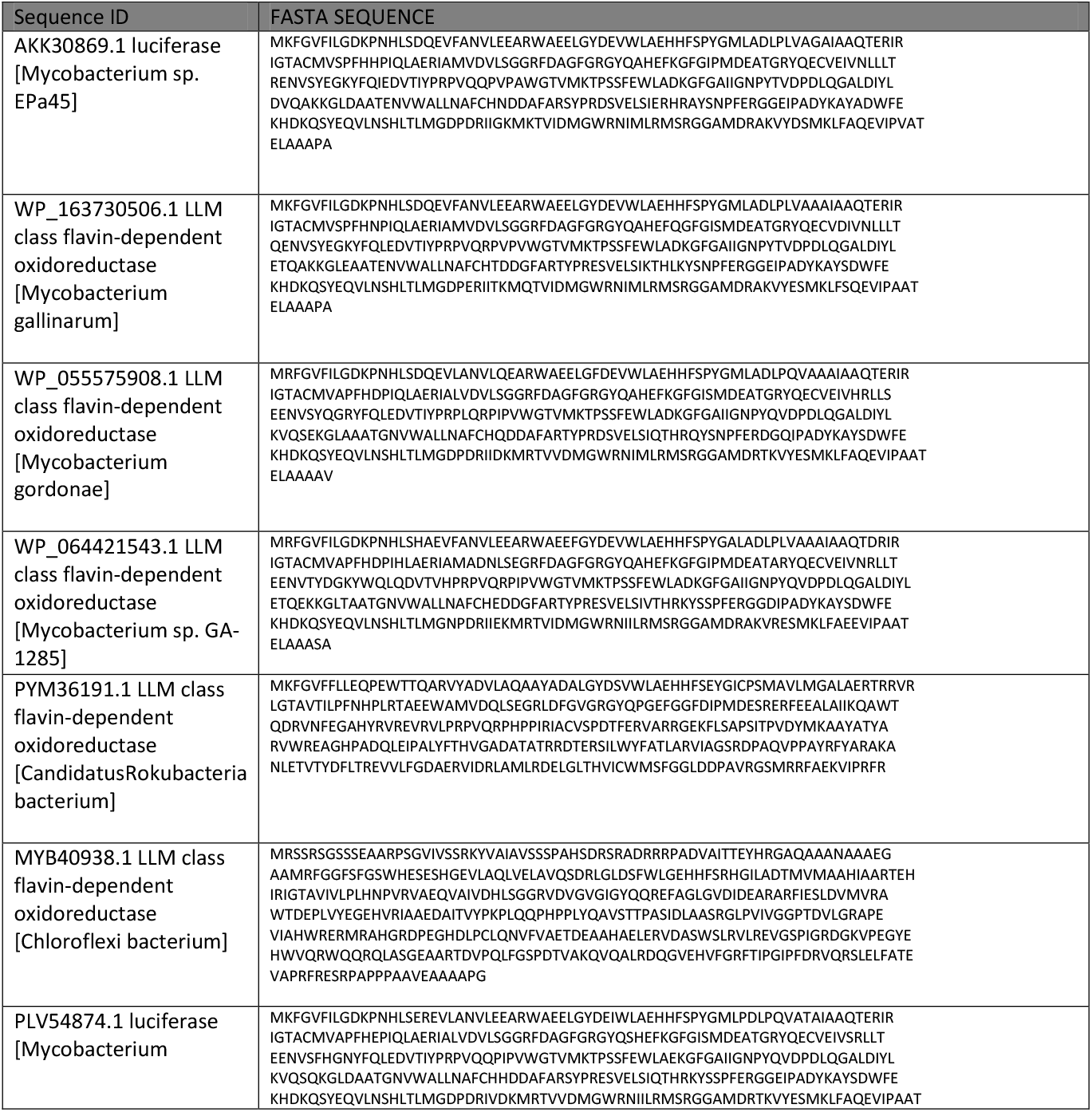
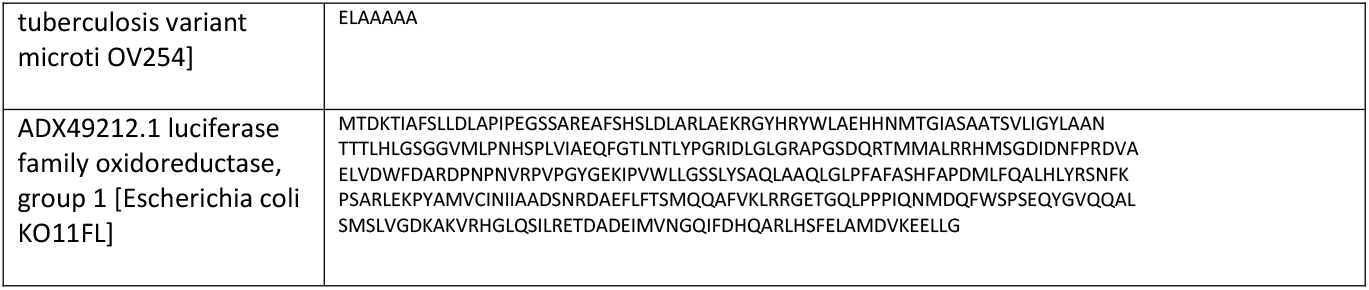
Luciferase and its related protein with their respective accession number and amino acid sequences.

### 2.2. Identification of Similar Proteins

The accession number was obtained from NCBI and was used as the reference sequence. BlastP was used to find proteins with similar sequences to AKK30869.1 luciferase for further analysis [https://blast.ncbi.nlm.nih.gov/Blast.cgi]^8^. The percentage similarity between the sequences was an essential parameter for consideration.

### 2.3 Studies on Multiple Sequence Alignment, Phylogenetic Trees and Guide Trees

The sequences of the Luciferase as well as those of the related proteins were aligned using CLUSTAL Omega tool, a multiple sequence alignment tool [https://www.ebi.ac.uk/Tools/msa/clustalo/]^9^. The evolutionary history was inferred by using the Maximum Likelihood method and Jones–Thornton– Taylor (JTT) matrix-based model in the MEGA software [https://www.megasoftware.net/]^10^. The percentage of trees in which the associated taxa clustered together is shown next to the branches. Initial tree(s) for the heuristic search were obtained automatically by applying Neighbor-Joining and BioNJ algorithms to a matrix of pairwise distances estimated using the JTT model, and then selecting the topology with superior log likelihood value. The tree is drawn to scale, with branch lengths measured in the number of substitutions per site.

### 2.4 Primary sequence analysis and physico-chemical characteristics

The amino acid sequence includes the signature from transcription and translation of gene and therefore bears various important information such as amino acid composition, physiochemical properties such as isoelectric point (pI), molecular weight (Mw), extinction coefficient (EC – quantitative study of protein – protein and protein – ligand interactions), instability index (II – stability of proteins), aliphatic index (AI – relative volume of protein occupied by aliphatic side chains), and Grand Average of Hydropathicities (GRAVY – sum of all hydropathicity values of all amino acids divided by number of residues in a sequence). Protparam was used to analyze and identify the physical and chemical properties of the protein in question [https://web.expasy.org/protparam/]^11^. A full list of characteristics identified can be found in the data section.

### 2.5 Secondary Structure analysis

The secondary structure of a protein refers to the arrangement of amino acid sequences in a polypeptide chain. So, the helix, sheet, and turn of amino acid sequence *Mycobacterium sp. EPa45* luciferase was predicted by PSI-blast based secondary structure PREDiction (PSIPRED) [http://bioinf.cs.ucl.ac.uk/psipred/]^12^.

### 2.6. Subcellular Localization

CELLO was used to detect the subcellular localization of the protein using a support vector mechanism (SVM) system [http://cello.life.nctu.edu.tw]^13^.

### 2.6 Peptide Cleavage Sites

PeptideCutter was used to identify the peptide cleavage sites of the protein [https://www.expasy.org/resources/peptidecutter]. Enzymes used for this theoretical identification include, but are not limited to: Arg-C Proteinase, CNBr, Iodosobenzoic acid and Proteinase K^14^. A full list of enzymes used can be found in the results section.

### 2.7 Prediction of Transmembrane helices

In order to make a prediction of membrane-spanning regions and their orientation in the *Mycobacterium sp. EPa45* luciferase, the TMPred software was used [https://embnet.vital-it.ch/software/TMPRED_form.html].

### 2.8 Conserved Domains Identification

Conserved domains as identifiers for the relationship between this protein and others like it were found using the NCBI Conserved Domain Database (CDD) program [https://www.ncbi.nlm.nih.gov/Structure/cdd/cdd.shtml]^15^. This was also correlated with the results obtained from MotifFinder software [https://www.genome.jp/tools/motif/].

### 2.9 Structural Visualization

Since there is no current 3D structure available for the *Mycobacterium sp. EPa45* luciferase I-TASSER, a program by the Zhang Lab at the University of Michigan, was used to determine the build the 3D structural model for the protein [https://zhanglab.ccmb.med.umich.edu/I-TASSER/]^16,17^. It also predicted structural qualities of each protein. Predicted qualities include: secondary structure, solvent accessibility, normalized B-factor and gene ontology. Visualizations of each protein were produced as well.^16,17^

### 2.10 Validation of structure obtained

RAMPAGE Ramachandran plot analysis was used for verification of 3D structures. It provides the number of residues in the favored, allowed, and outlier region. If a good proportion of residues lie in the favored and allowed region, then the model is predicted to be good. The quality of the models were also accessed using PROSA, PROCHECK and Verify 3D [https://prosa.services.came.sbg.ac.at/prosa.php] [https://servicesn.mbi.ucla.edu/PROCHECK/] [https://servicesn.mbi.ucla.edu/Verify3D/]^18–20^. The stereochemical quality of the submitted models based on its phi/psi angle arrangement was analyzed by PROCHECK and RAMPAGE and then Ramachandran plots were generated which highlights the percentage of residues in the favored, allowed or in outlier regions. The model is considered to be good if a greater proportion of the residues lie in the favored and allowed region. ProSa on the other hand does a comparative analysis by calculating the potential energy of the protein models and comparing them to the experimental structures deposited in the PDB. The Z-Scores obtained from each model suggest that the structures are comparable to the NMR structures of similar size. The local quality of the protein model was evaluated by Verify3D on the basis of structure-sequence compatibility to generate a compatibility value for each residue of the protein. A model with 80% of their residues with a 3D-1D score equal to or higher than 0.2 is considered to be a high quality structure.

## 3. Results

### 3.1 Sequence retrieval and phylogenetic analysis

The accession number AKK30869.1 for *Mycobacterium sp. EPa45 luciferase* was obtained from NCBI and was used as the reference sequence. Identification of similar proteins was done via BLASTP. These proteins along with the *Mycobacterium sp. EPa45* luciferase are listed in Table 1. The phylogenetic tree was constructed using Maximum Likelihood method and JTT matrix-based model. The tree with the highest log likelihood (−4434.19) is shown. The phylogenetic tree suggested a close similarity between *Mycobacterium sp. EPa45 luciferase* and *Mycobacterium gallinarum* LLM class flavin-dependent oxidoreductase. On the other hand however, it was farther from the *Escherichia coli KO11FL* ADX49212.1 luciferase family oxidoreductase, group 1. The phylogenetic tree is shown in Figure 1.

**Figure 1:**
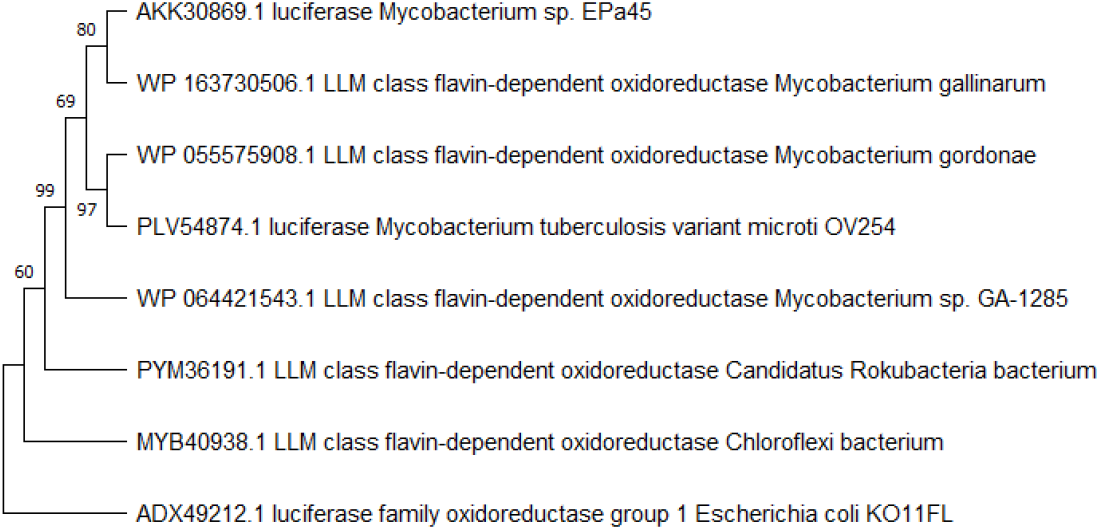
Phylogenetic Analysis of Luciferase and related protein. The evolutionary history was inferred by using the Maximum Likelihood method and JTT matrix-based model

### 3.2 Physiochemical Characterization

The primary purpose of comparing our protein of interest with those of luciferases in other pathogenic and non-pathogenic bacteria, as shown in Table 1, was to see which ones of the two it bore any similarity to, if at all. The Protparam analysis performed, shown in Table 2, exhibits that there are quite a few physiochemical similarities between the luciferase in *Mycobacterium sp. EPa45, Mycobacterium Tuberculosis* and *Mycobacterium gallinarum*. These strains have three hundred and fifty-seven amino acids in their chains. Additionally, their number of charged residues, along with instability index, aliphatic indexes are extremely similar. Comparatively, the luciferase found in *Escherichia coli* is shorter and shows a lesser degree of physiochemical similarities. (18) A detailed description of the physiochemical properties of luciferase and its related proteins can be found in Table 2.

**Table 2:** A table of physiochemical properties as analyzed by Expasy’s Protparam program

| Protein | # Amino Acids | Molecular Weight | pI | -R | +R | Extinction Coefficient | Instability Index | Aliphatic Index | GRA VY |
| --- | --- | --- | --- | --- | --- | --- | --- | --- | --- |
| AKK30869.1 luciferase [Mycobacterium sp. EPa45] | 357 | 40063.49 | 5.02 | 51 | 34 | 60975, 60850 | 38.58 | 81.48 | - 0.237 |
| WP_163730506.1 LLM class flavin-dependent oxidoreductase [Mycobacterium gallinarum] | 357 | 40123.53 | 4.93 | 50 | 32 | 60975, 60850 | 33.64 | 80.92 | - 0.240 |
| WP_055575908.1 LLM class flavin-dependent oxidoreductase [Mycobacterium gordonae] | 357 | 40272.64 | 5.12 | 49 | 34 | 59485, 59360 | 32.27 | 82.30 | - 0.265 |
| WP_064421543.1 LLM class flavin-dependent oxidoreductase [Mycobacterium sp. GA-1285] | 357 | 40200.47 | 5.21 | 52 | 36 | 63495, 63370 | 34.29 | 79.33 | - 0.303 |
| PYM36191.1 LLM class flavin-dependent oxidoreductase [CandidatusRokubacteria bacterium] | 349 | 39726.39 | 6.73 | 45 | 44 | 59485, 59360 | 41.20 | 81.40 | - 0.205 |
| MYB40938.1 LLM class flavin-dependent oxidoreductase [Chloroflexi bacterium] | 444 | 48211.11 | 5.95 | 57 | 46 | 48930 | 44.01 | 84.41 | - 0.254 |
| PLV54874.1 luciferase [Mycobacterium tuberculosis variant microti OV254] | 357 | 40198.53 | 5.15 | 50 | 34 | 59485, 59360 | 38.53 | 81.46 | - 0.274 |
| ADX49212.1 luciferase family oxidoreductase, group 1 [Escherichia coli KO11FL] | 335 | 37106.30 | 5.93 | 36 | 29 | 35410 | 35.81 | 87.46 | - 0.167 |

When looking at the CLUSTL alignment of the three sequences (Figure 2), it becomes quite clear why these similarities arise. There are a great number of conserved sequences between the luciferase in *Mycobacterium sp. EPa45* and *Mycobacterium Tuberculosis*. This is not the case with *Escherichia coli*. Additionally, the phylogram generated clearly demonstrates that *Mycobacterium sp. EPa45* and *Mycobacterium Tuberculosis* are more closely related to each other than either are to *Escherichia coli*. Given these observations, our luciferase of interest, that of *Mycobacterium sp. EPa45*, can be said to be closer in structure and characteristics to pathogenic bacterium instead of non-pathogenic bacterium.

**Figure 2:**
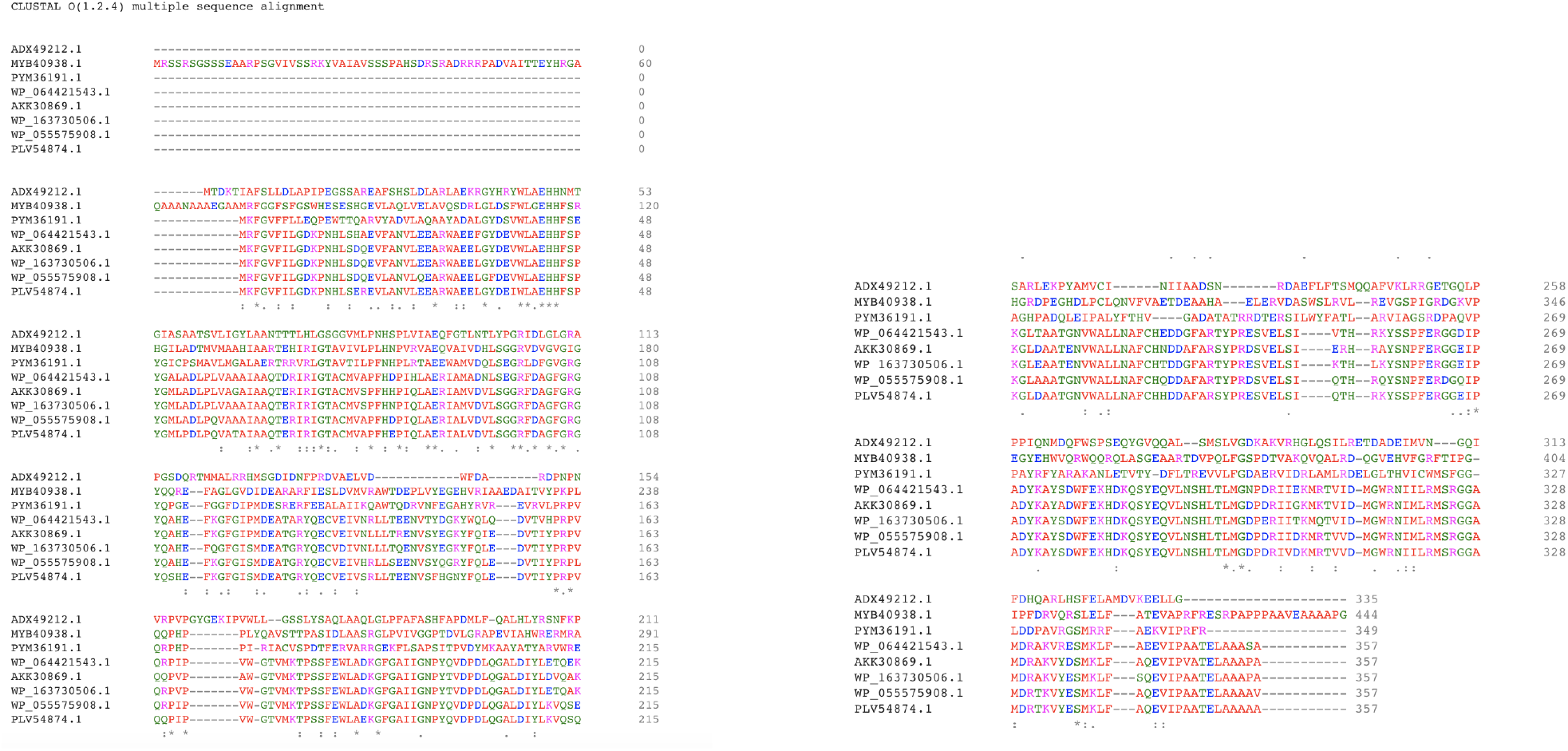
Multiple Sequence Alignment of Luciferase and related proteins

### 3.3 Protein Secondary Structure analysis

A detailed analysis into the secondary structure of bacterial luciferase and its related proteins were done and are presented in table 3. The distribution percentage of *Mycobacterium sp. EPa45 luciferase* secondary structure was 48.74% alpha helix, 4.20% beta turn, 13.74% extended strand and 33.33% random coil [Figure 3]. Prediction of transmembrane helices and their topology can be found in Figure 4A. Schematic representation of protein domain assignment from the secondary structure of *Mycobacterium sp. EPa45 luciferase* has been provided in Figure 4B.

**Figure 3:**
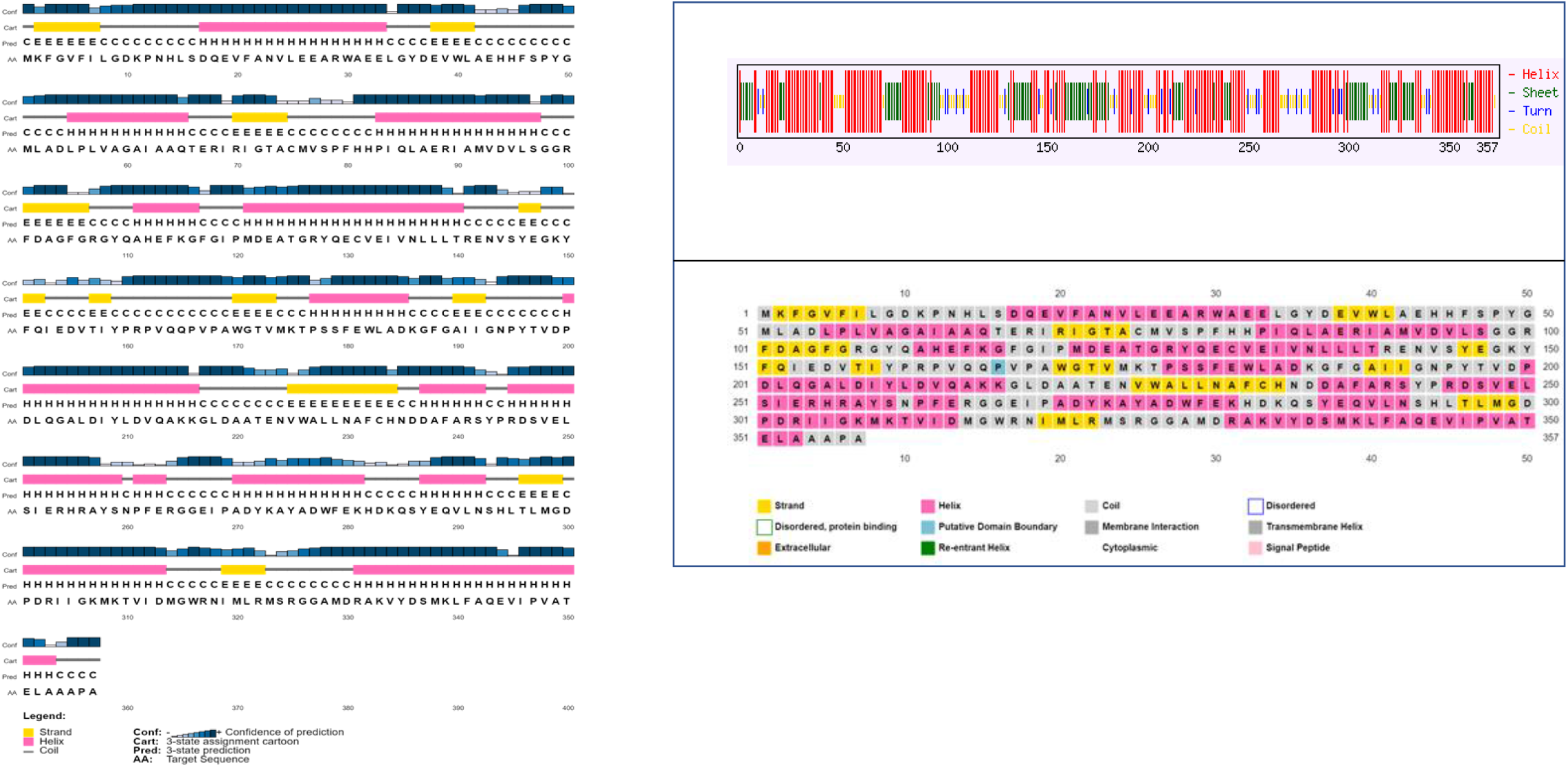
Secondary structure analysis of AKK30869.1 luciferase [*Mycobacterium sp. EPa45*]

**Figure 4:**
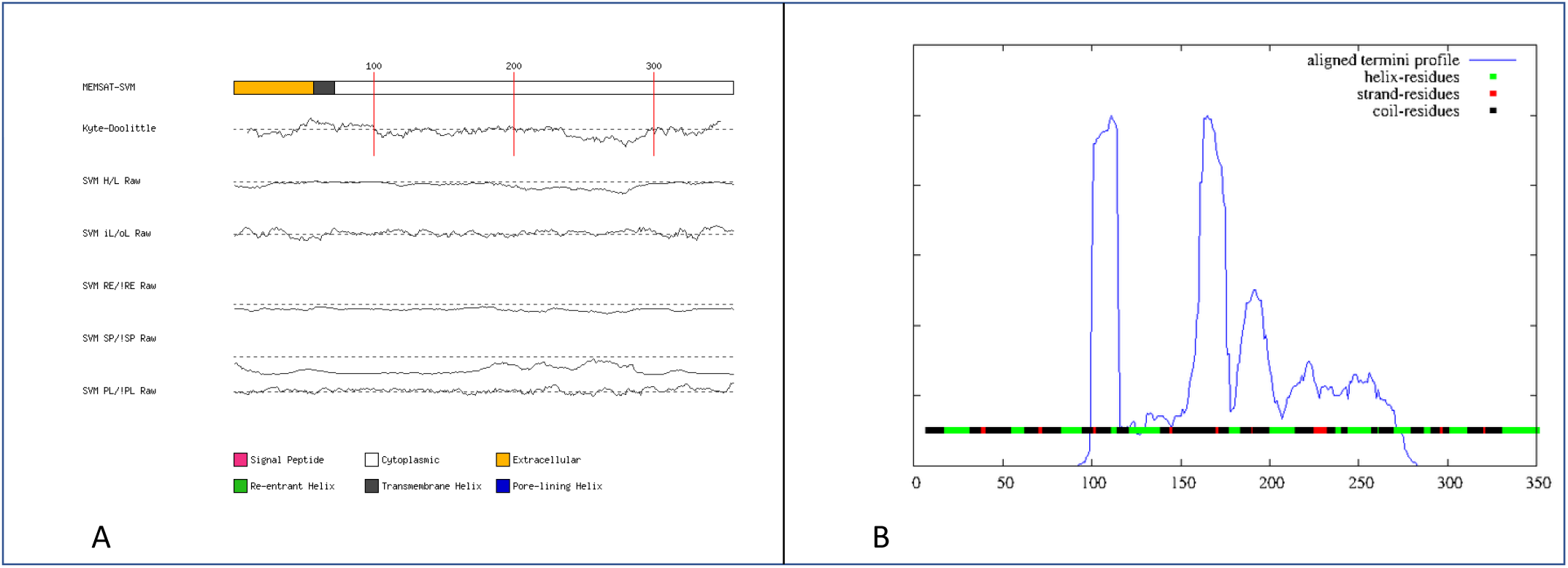
**A**. Prediction of trans membrane helices and their topology **B**. Protein domain assignment from secondary structure of AKK30869.1 luciferase [*Mycobacterium sp. EPa45*]

**Table 3:** A table of analysis of protein secondary structure done using SOPMA

| Protein | # Amino Acids | Alpha Helix | Beta Turn | Extended Strand | Random Coil |
| --- | --- | --- | --- | --- | --- |
| AKK30869.1 luciferase [Mycobacterium sp. EPa45] | 357 | 174, 48.74% | 15, 4.20% | 49, 13.74% | 119, 33.33% |
| WP_163730506.1 LLM class flavin-dependent oxidoreductase [Mycobacterium gallinarum] | 357 | 116, 46.50% | 18, 5.04% | 58, 16.25% | 115, 32.21% |
| WP_055575908.1 LLM class flavin-dependent oxidoreductase [Mycobacterium gordonae] | 357 | 174, 48.74% | 20, 5.60% | 55, 15.41% | 108, 30.25% |
| WP_064421543.1 LLM class flavin-dependent oxidoreductase [Mycobacterium sp. GA-1285] | 357 | 173, 48.46% | 17, 4.76% | 53, 14.85% | 114, 31.93% |
| PYM36191.1 LLM class flavin-dependent oxidoreductase [CandidatusRokubacteria bacterium] | 349 | 160, 45.85% | 15, 4.30% | 60, 17.19% | 114, 32.66% |
| MYB40938.1 LLM class flavin-dependent oxidoreductase [Chloroflexi bacterium] | 444 | 175, 39.41% | 21, 4.73% | 73, 16.44% | 175, 39.41% |
| PLV54874.1 luciferase [Mycobacterium tuberculosis variant microti OV254] | 357 | 181, 50.70% | 18, 5.04% | 53, 14.85% | 105, 29.41% |
| ADX49212.1 luciferase family oxidoreductase, group 1 [Escherichia coli KO11FL] | 335 | 160, 47.76% | 15, 4.48% | 51, 15.22% | 109, 32.54% |

### 3.4 Functional analysis of the Protein

The prediction of transmembrane helices is shown in Figure 5. The conserved domain database search tells us that the AKK30869.1 luciferase protein falls under the superfamily of flavin utilizing monooxygenases, indicative of the fact that this protein is an oxidative luciferase which is meant to aid in fluorescence. The purpose of flavin utilizing monooxygenases is quite different from this. They are tasked with eliminating insoluble compounds in organisms. This process is known as xenobiotics. Generally, these proteins are meant to interact with FAD molecules. (1) In this particular case, according to the CDD search, this monooxygenase also falls into the subgroup of luciferase-like monooxygenases (LLMs). It is important to note that not all monooxygenases act as luciferases, however, most bacterial luciferases have a catalyzed mechanism which includes an oxidative process. (2) The findings from the CDD analysis correlated with that of the Motiffinder which suggested the presence of 2 motifs Nitrosopumilus output domain 5 and Luciferase-like monooxygenase [Figure 6]. Table 4 gives the prediction of subcellular localization of *Mycobacterium sp. EPa45 luciferase*. Peptide cleavage site analysis by different enzymes for *Mycobacterium sp. EPa45 luciferase* is listed in Table 5.

**Figure 5:**
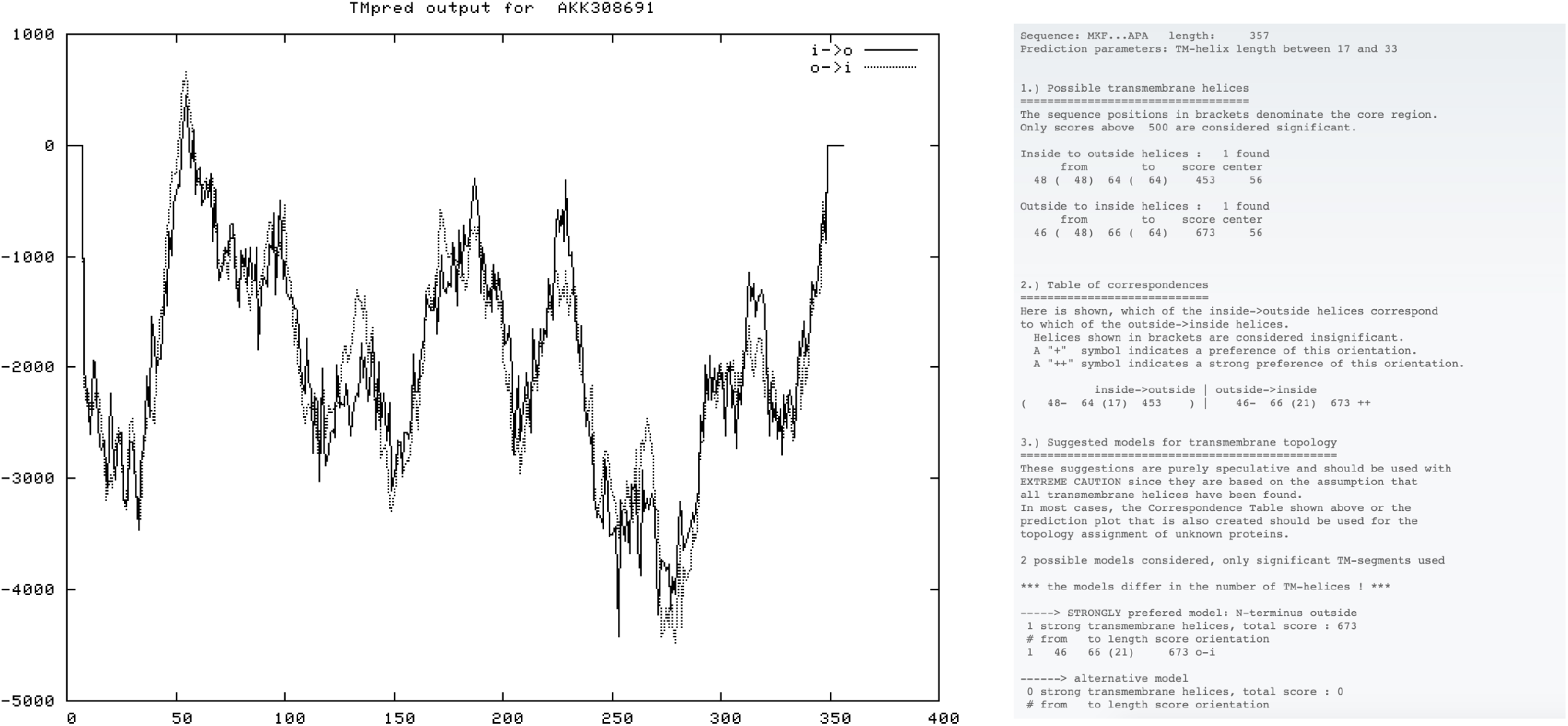
Prediction of transmembrane helices and orientation in AKK30869.1 luciferase [*Mycobacterium sp. EPa45*]

**Figure 6:**
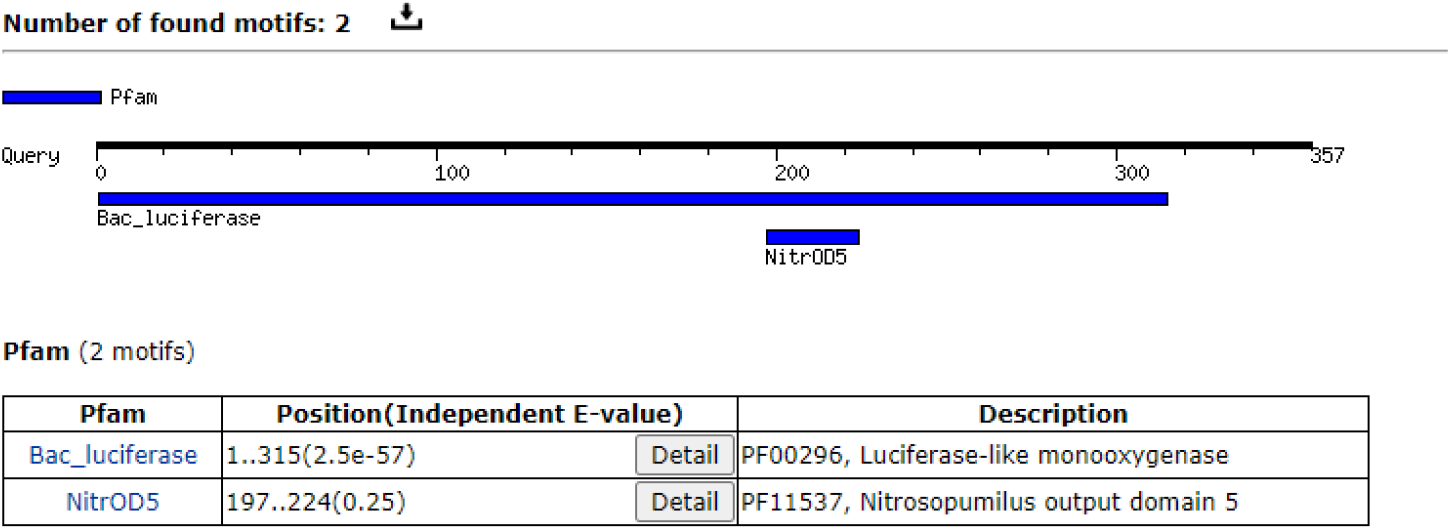
Result of motif finder showing two functional motifs for the AKK30869.1 luciferase [*Mycobacterium sp. EPa45*]

**Table 4:** Analysis of subcellular localization of protein by CELLO

|  |  |  |
| --- | --- | --- |
| SeqID: AKK30869.1 luciferase [Mycobacterium sp. EPa45] |  |  |
| Analysis Report: |  |  |
| SVM | LOCALIZATION | RELIABILITY |
| Amino Acid Comp. | Cytoplasmic | 0.933 |
| N-peptide Comp. | Cytoplasmic | 0.827 |
| Partitioned seq. Comp. | Cytoplasmic | 0.943 |
| Physico-chemical Comp. | Membrane | 0.468 |
| Neighboring seq. Comp. | Cytoplasmic | 0.986 |
| CELLO Prediction: |  |  |
|  | Cytoplasmic | 4.126 * |
|  | Membrane | 0.711 |
|  | Extracellular | 0.153 |
|  | CellWall | 0.011 |
| ***** |  |  |

**Table 5:**
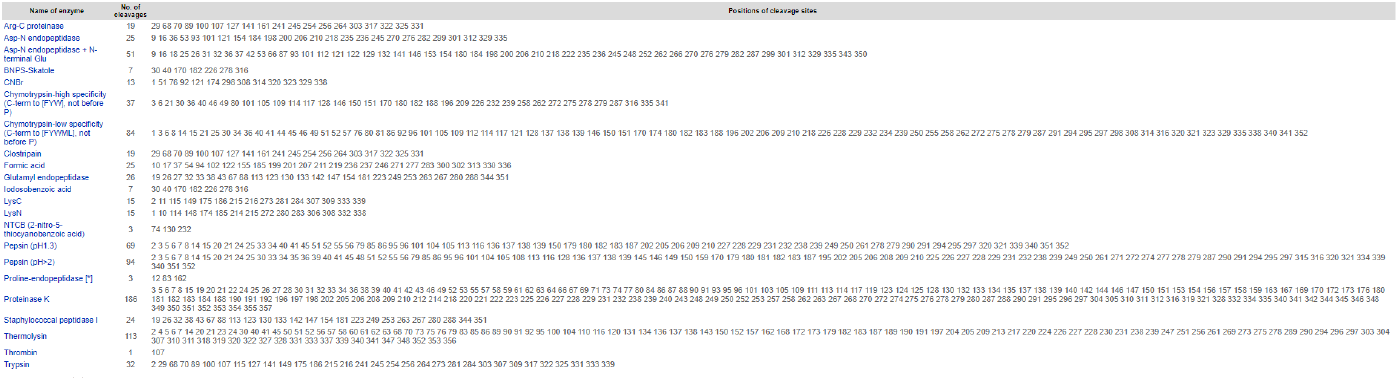
Peptide cleavage site analysis by PeptideCutter

### 3.5 3D protein modeling and its validation

I-TASSER web server enabled generation of the 3D structure of the protein [Figure 7]. It provides five models based on the amino acid sequence, and each model is assigned an individual confidence scores (C-score) calculated based on the significance of threading template alignments and the convergence parameters of the structure assembly simulations. For each of the proteins the models with a higher C-score was chosen and was subjected to further analysis by RAMPAGE, PROSA, PROCHECK and VERIFY3D. All the models chosen for the respective proteins were found to be of better quality in terms of stereochemical quality and potential energy. The Ramachandran plots obtained for the respective protein can be found in Figure 8–9. The analysis by PROSA and VERIFY3D can be found in Figure 10–17.

**Figure 7:**
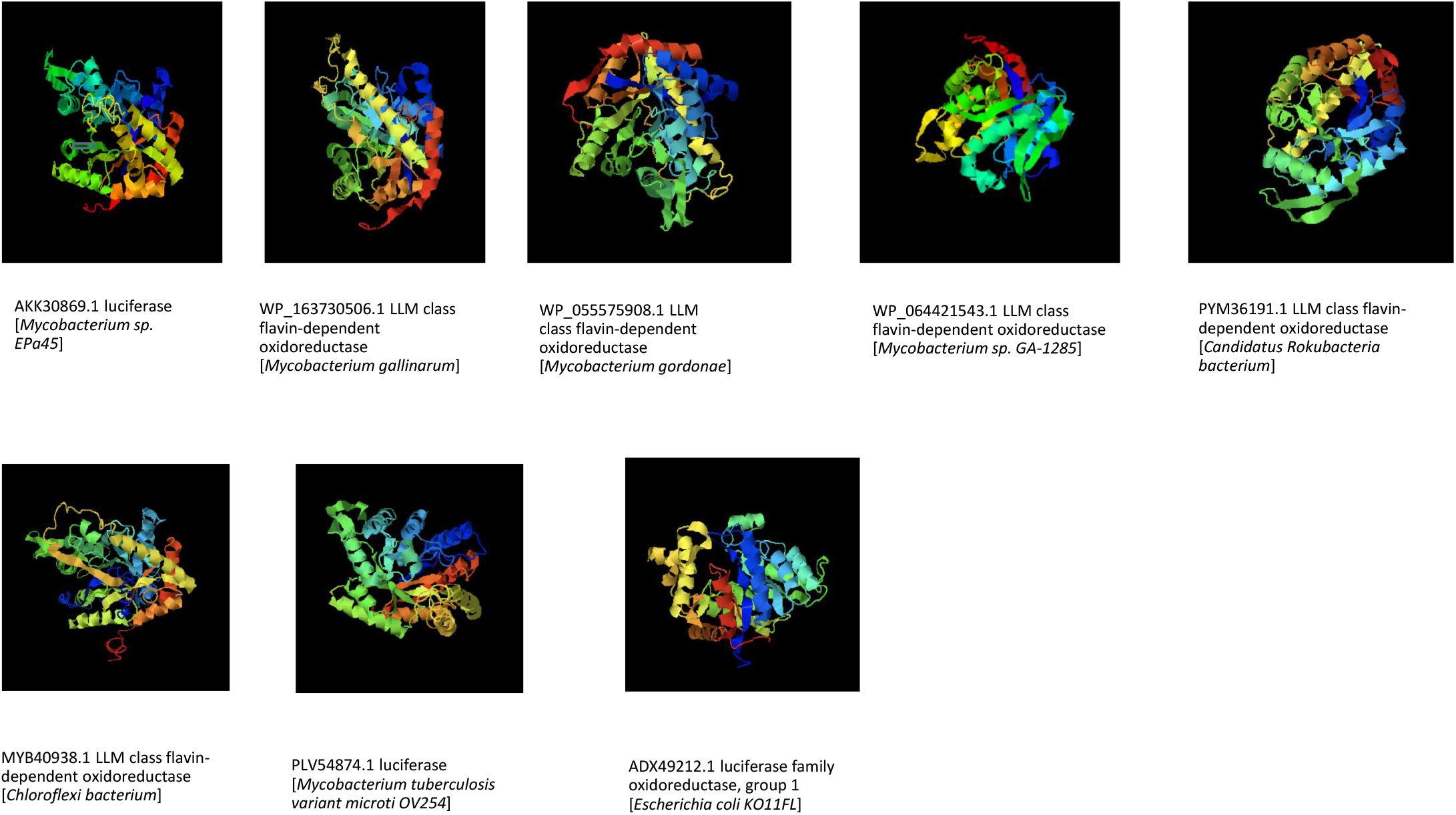
3D structure of luciferase and its related proteins generated by I-TASSER

**Figure 8:**
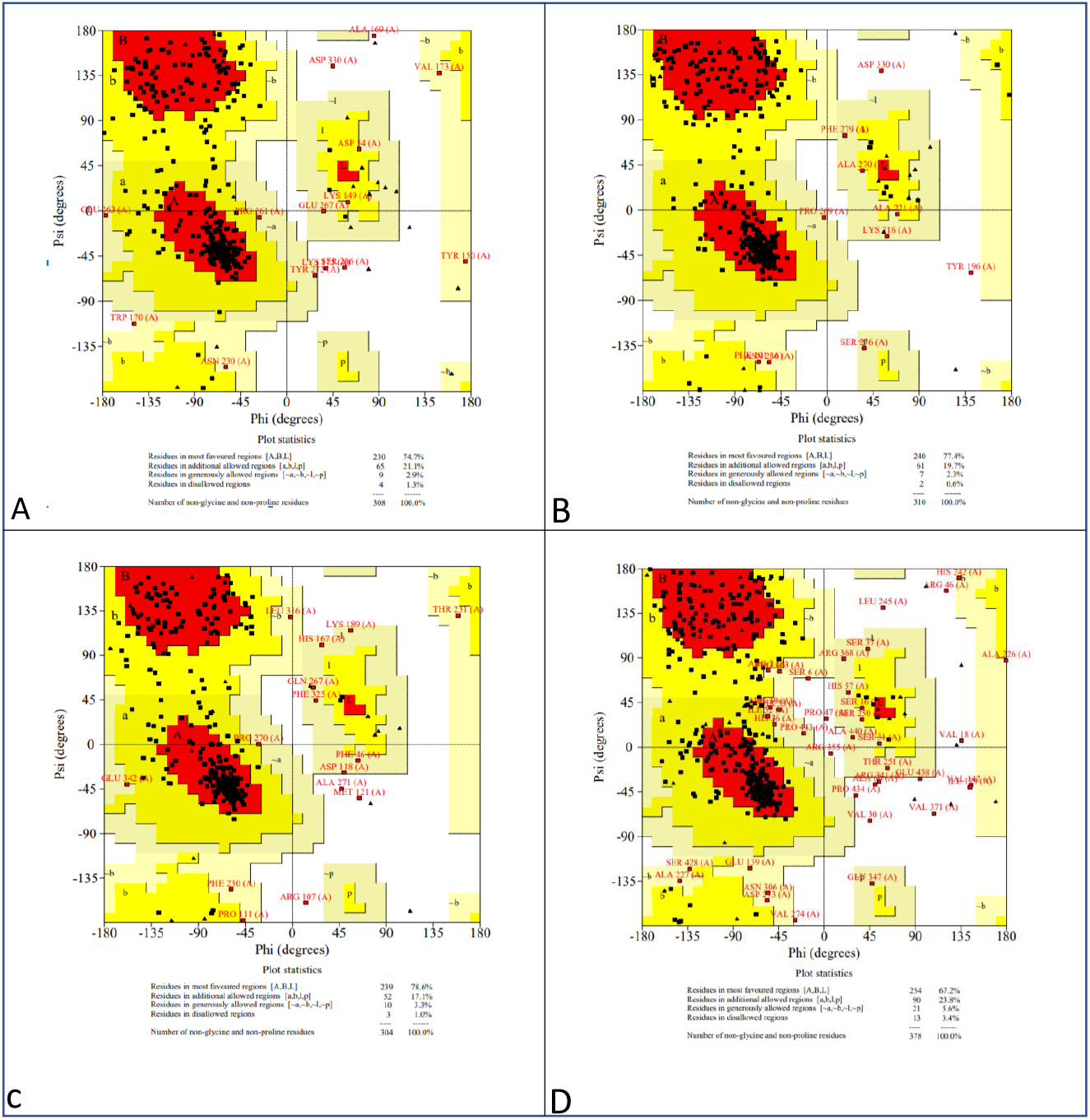
Ramachandran Plot of a. AKK30869.1 luciferase [*Mycobacterium sp. EPa45*] b. WP_163730506.1 LLM class flavin-dependent oxidoreductase [*Mycobacterium gallinarum*] c. PYM36191.1 LLM class flavin-dependent oxidoreductase [*Candidatus Rokubacteria bacterium*] d. MYB40938.1 LLM class flavin-dependent oxidoreductase [*Chloroflexi bacterium*]

**Figure 9:**
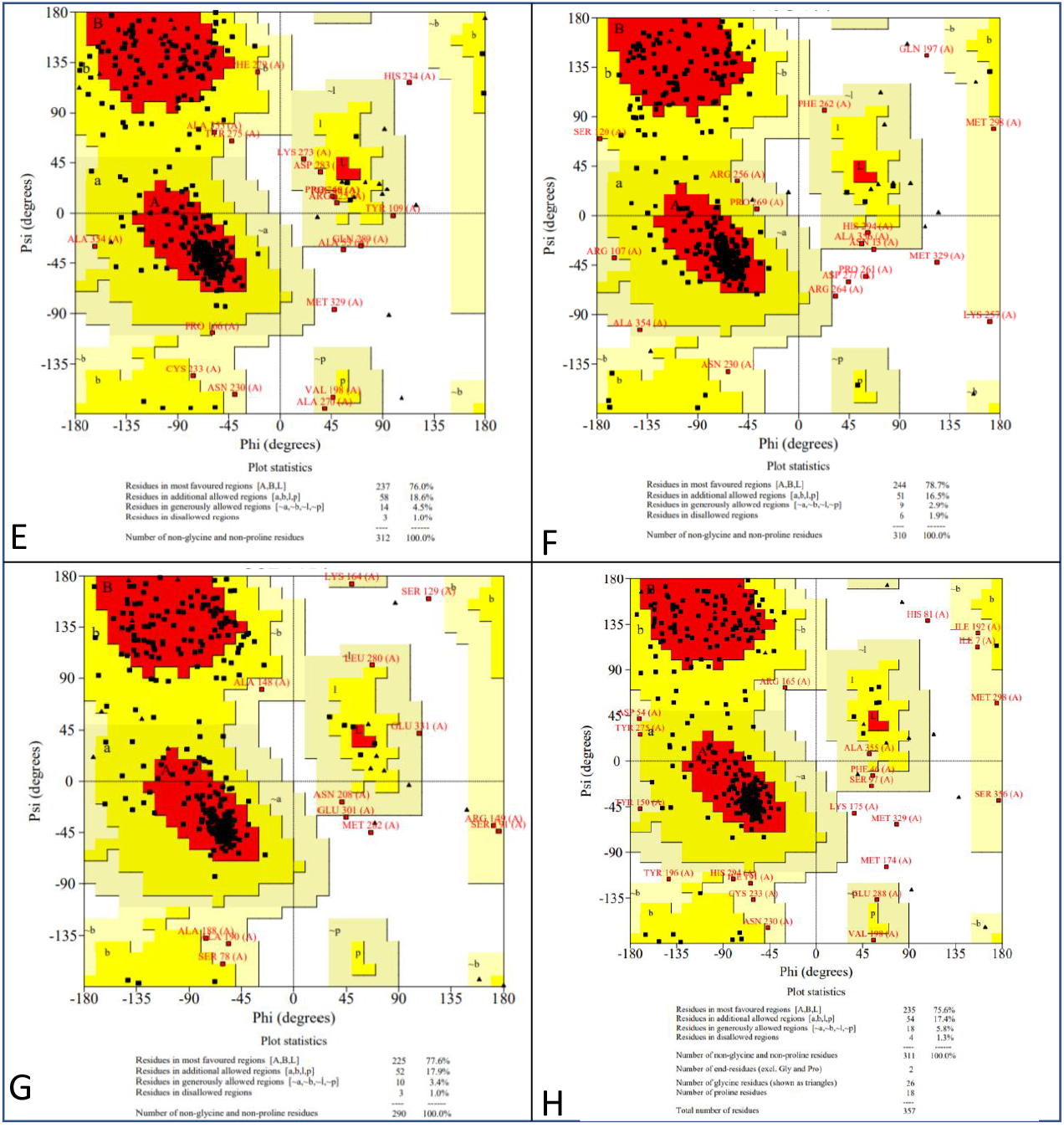
Ramachandran Plot of e. WP_055575908.1 LLM class flavin-dependent oxidoreductase [*Mycobacterium gordonae*] f. PLV54874.1 luciferase [*Mycobacterium tuberculosis* variant microti OV254] g. ADX49212.1 luciferase family oxidoreductase, group 1 [*Escherichia coli* KO11FL] h. WP_064421543.1 LLM class flavin-dependent oxidoreductase [*Mycobacterium sp*. GA-1285]

**Figure 10:**
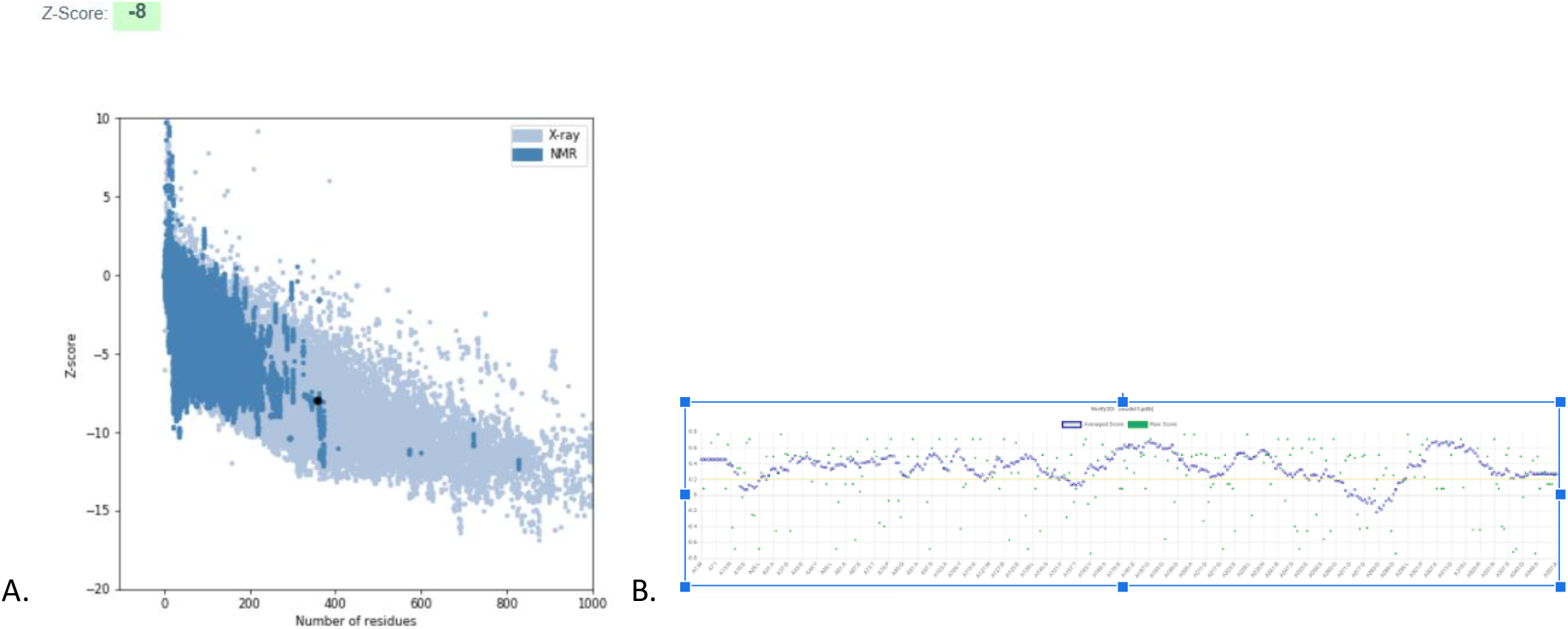
Validation of the in silico predicted model of AKK30869.1 luciferase [*Mycobacterium sp. EPa45*] by ProSa and Verify3D. A. Structure validation by ProSa, which shows the Z-score of the predicted model of Nucleoprotein (black dot), when compared to a non-redundant set of crystallographic structures (light blue dots) and NMR structures (dark blue dots). B. Structure validation by Verify3D, which shows the 3D-1D score for each atom of the predicted model of Nucleoprotein. The graphic shows that 86.55% of the residues of the in silico structure of Nucleoprotein presented a compatibility score of 0.2 or higher, which indicates that the structure is a high-quality model according to Verify3D.

**Figure 11:**
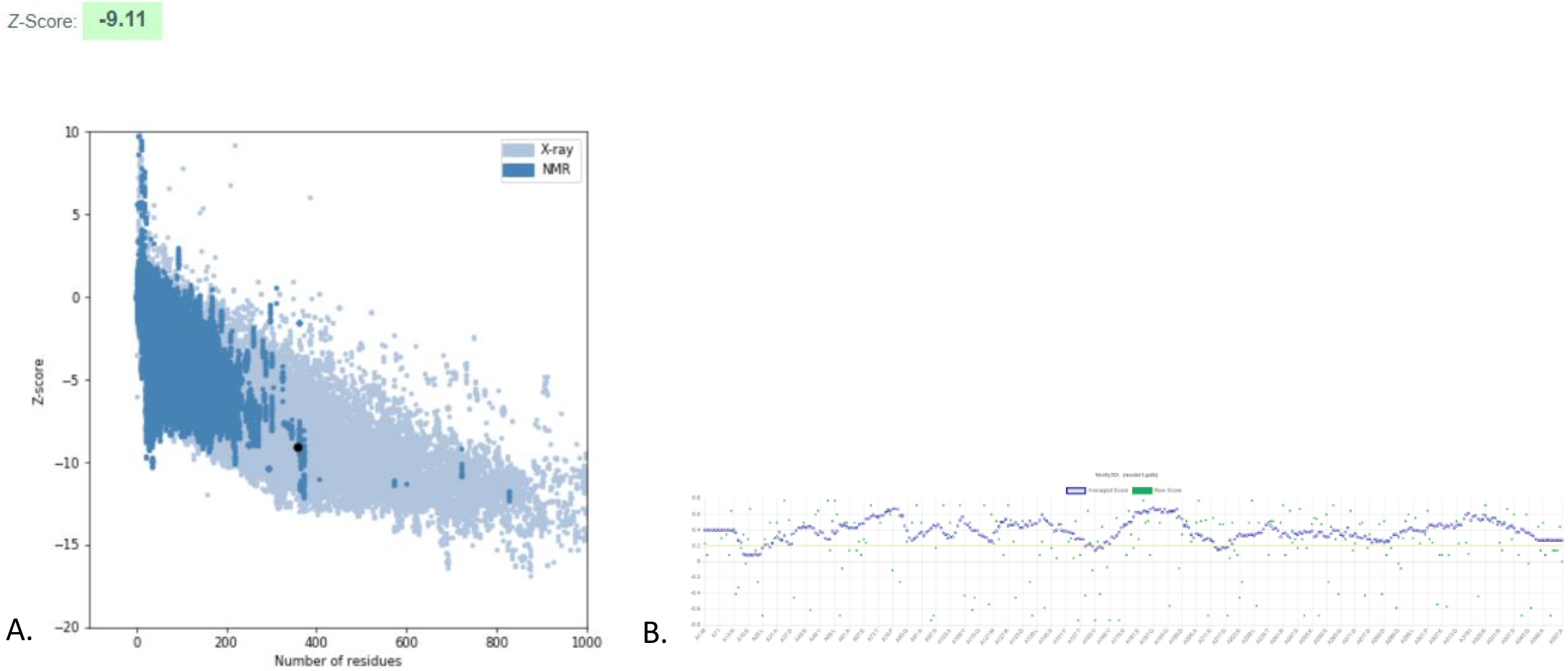
Validation of the in silico predicted model of WP_163730506.1 LLM class flavin-dependent oxidoreductase [*Mycobacterium gallinarum*] by ProSa and Verify3D. A. Structure validation by ProSa, which shows the Z-score of the predicted model of Nucleoprotein (black dot), when compared to a non-redundant set of crystallographic structures (light blue dots) and NMR structures (dark blue dots). B. Structure validation by Verify3D, showing the 3D-1D score for each atom of the predicted model of Nucleoprotein. The graphic shows 3.84% of the residues of the in silico structure of Nucleoprotein presented a compatibility score of 0.2 or higher, which indicates that the structure is a high-quality model according to Verify3D.

**Figure 12:**
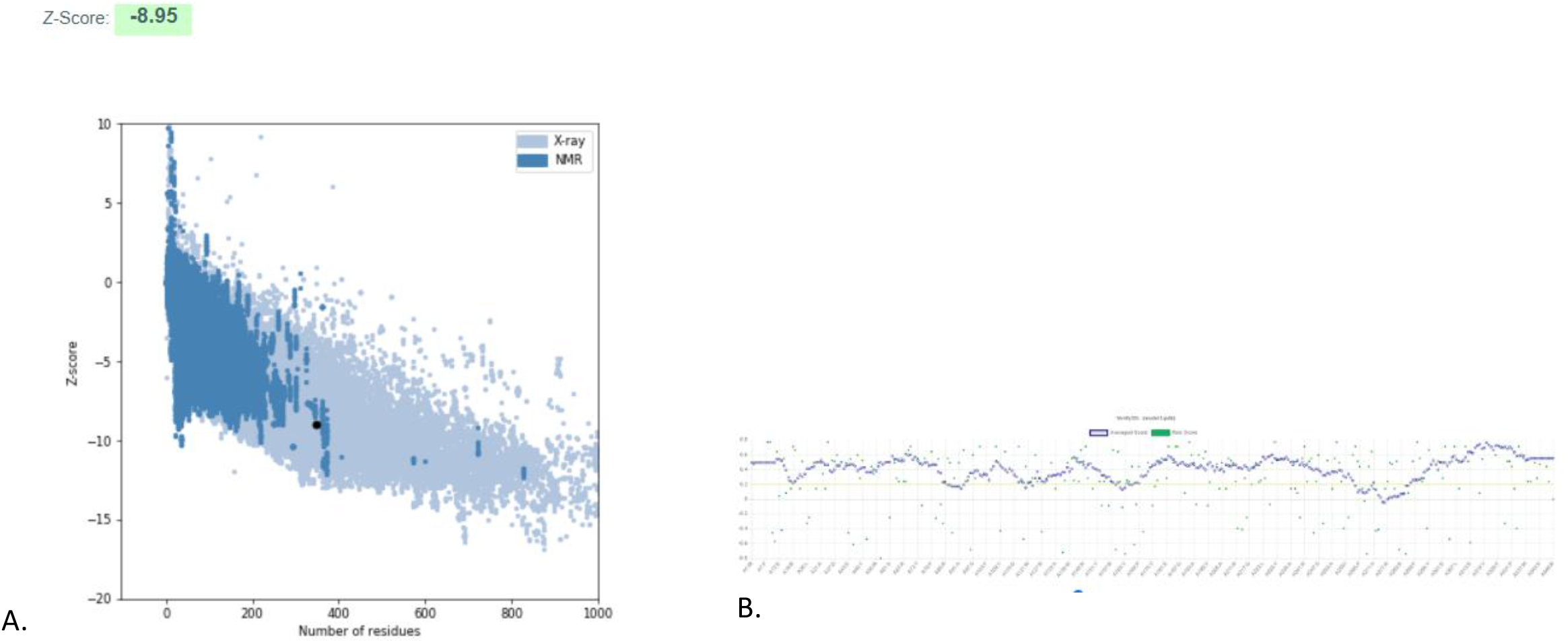
Validation of the in silico predicted model of PYM36191.1 LLM class flavin-dependent oxidoreductase [*CandidatusRokubacteria bacterium*] by ProSa and Verify3D. A. Structure validation by ProSa, which shows the Z-score of the predicted model of Nucleoprotein (black dot), when compared to a non-redundant set of crystallographic structures (light blue dots) and NMR structures (dark blue dots). B. Structure validation by Verify3D, which shows the 3D-1D score for each atom of the predicted model of Nucleoprotein. The graphic shows that 89.11% of the residues of the in silico structure of Nucleoprotein presented a compatibility score of 0.2 or higher, which indicates that the structure is a high-quality model according to Verify3D.

**Figure 13:**
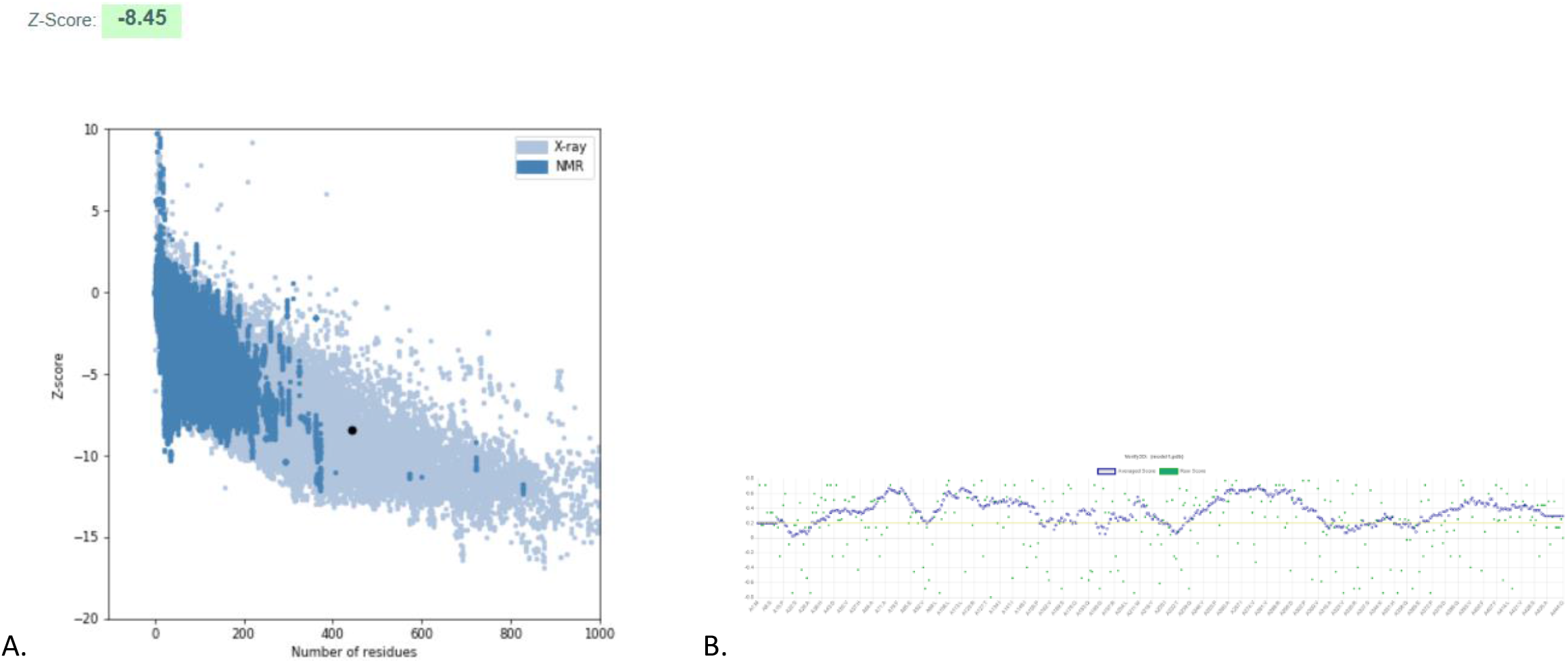
Validation of the in silico predicted model of MYB40938.1 LLM class flavin-dependent oxidoreductase [*Chloroflexi bacterium*] by ProSa and Verify3D. A. Structure validation by ProSa, which shows the Z-score of the predicted model of Nucleoprotein (black dot), when compared to a non-redundant set of crystallographic structures (light blue dots) and NMR structures (dark blue dots). B. Structure validation by Verify3D, which shows the 3D-1D score for each atom of the predicted model of Nucleoprotein. The graphic shows that 82.66% of the residues of the in silico structure of Nucleoprotein presented a compatibility score of 0.2 or higher, which indicates that the structure is a high-quality model according to Verify3D.

**Figure 14:**
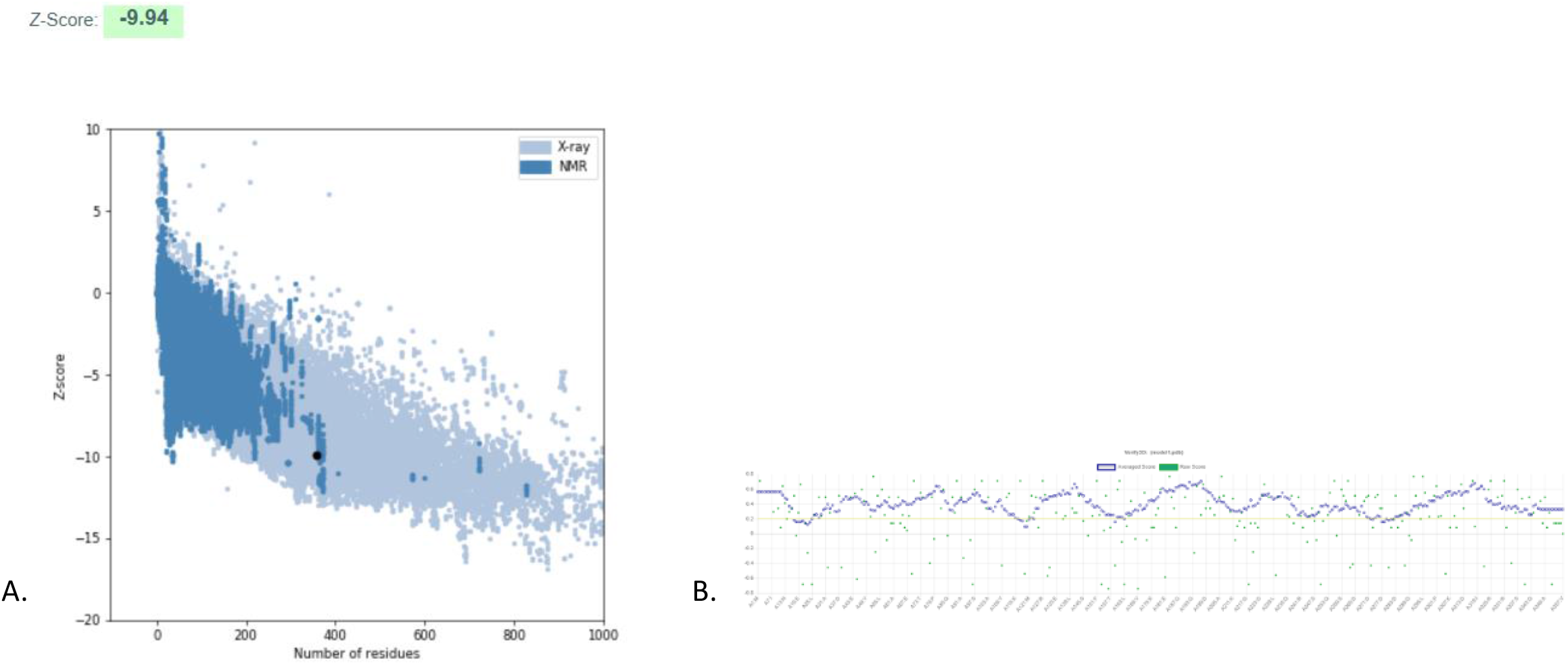
Validation of the in silico predicted model of WP_055575908.1 LLM class flavin-dependent oxidoreductase [*Mycobacterium gordonae*] by ProSa and Verify3D. A. Structure validation by ProSa, which shows the Z-score of the predicted model of Nucleoprotein (black dot), when compared to a non-redundant set of crystallographic structures (light blue dots) and NMR structures (dark blue dots). B. Structure validation by Verify3D, which shows the 3D-1D score for each atom of the predicted model of Nucleoprotein. The graphic shows that 94.4% of the residues of the in silico structure of Nucleoprotein presented a compatibility score of 0.2 or higher, which indicates that the structure is a high-quality model according to Verify3D.

**Figure 15:**
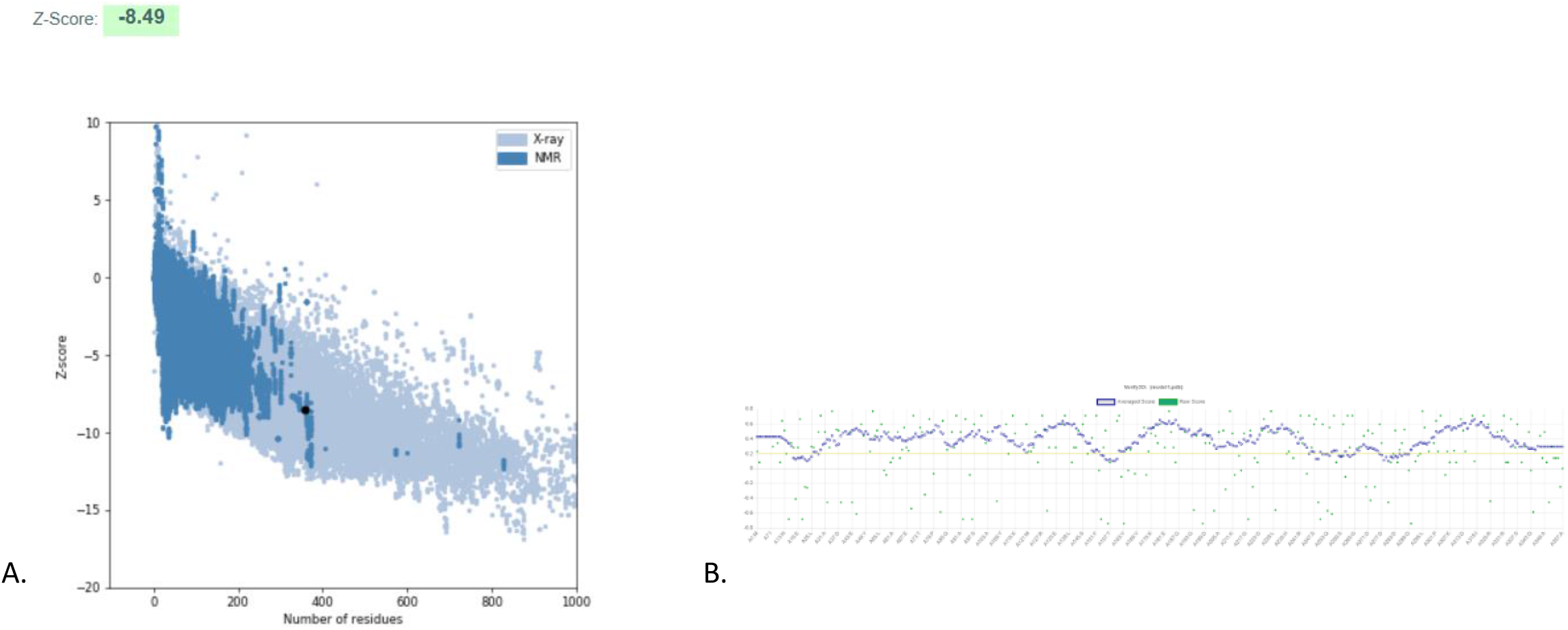
Validation of the in silico predicted model of PLV54874.1 luciferase [*Mycobacterium tuberculosis* variant microti OV254] by ProSa and Verify3D. A. Structure validation by ProSa, which shows the Z-score of the predicted model of Nucleoprotein (black dot), when compared to a non-redundant set of crystallographic structures (light blue dots) and NMR structures (dark blue dots). B. Structure validation by Verify3D, which shows the 3D-1D score for each atom of the predicted model of Nucleoprotein. The graphic shows that 88.52% of the residues of the in silico structure of Nucleoprotein presented a compatibility score of 0.2 or higher, which indicates that the structure is a high-quality model according to Verify3D.

**Figure 16:**
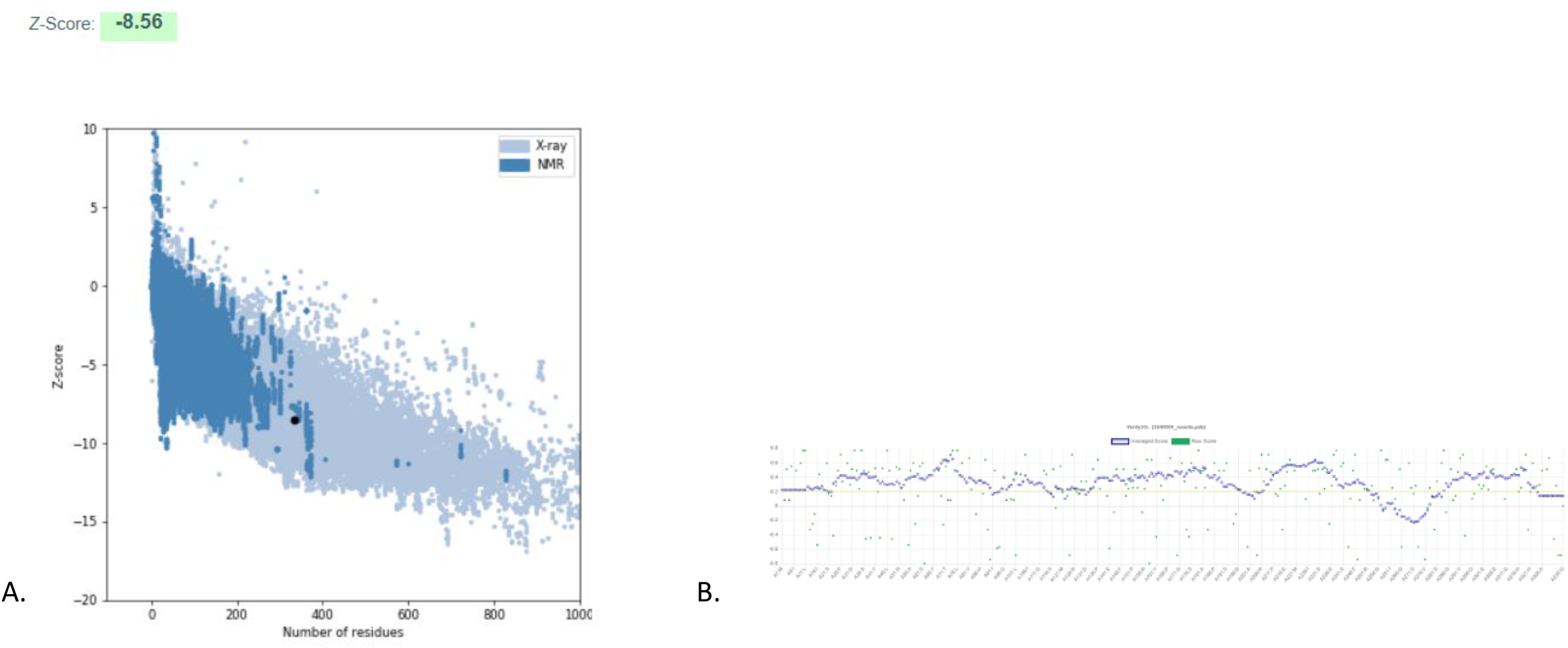
Validation of the in silico predicted model of ADX49212.1 luciferase family oxidoreductase, group 1 [*Escherichia coli* KO11FL] by ProSa and Verify3D. A. Structure validation by ProSa, which shows the Z-score of the predicted model of Nucleoprotein (black dot), when compared to a non-redundant set of crystallographic structures (light blue dots) and NMR structures (dark blue dots). B. Structure validation by Verify3D, which shows the 3D-1D score for each atom of the predicted model of Nucleoprotein. The graphic shows that 82.09% of the residues of the in silico structure of Nucleoprotein presented a compatibility score of 0.2 or higher, which indicates that the structure is a high-quality model according to Verify3D.

**Figure 17:**
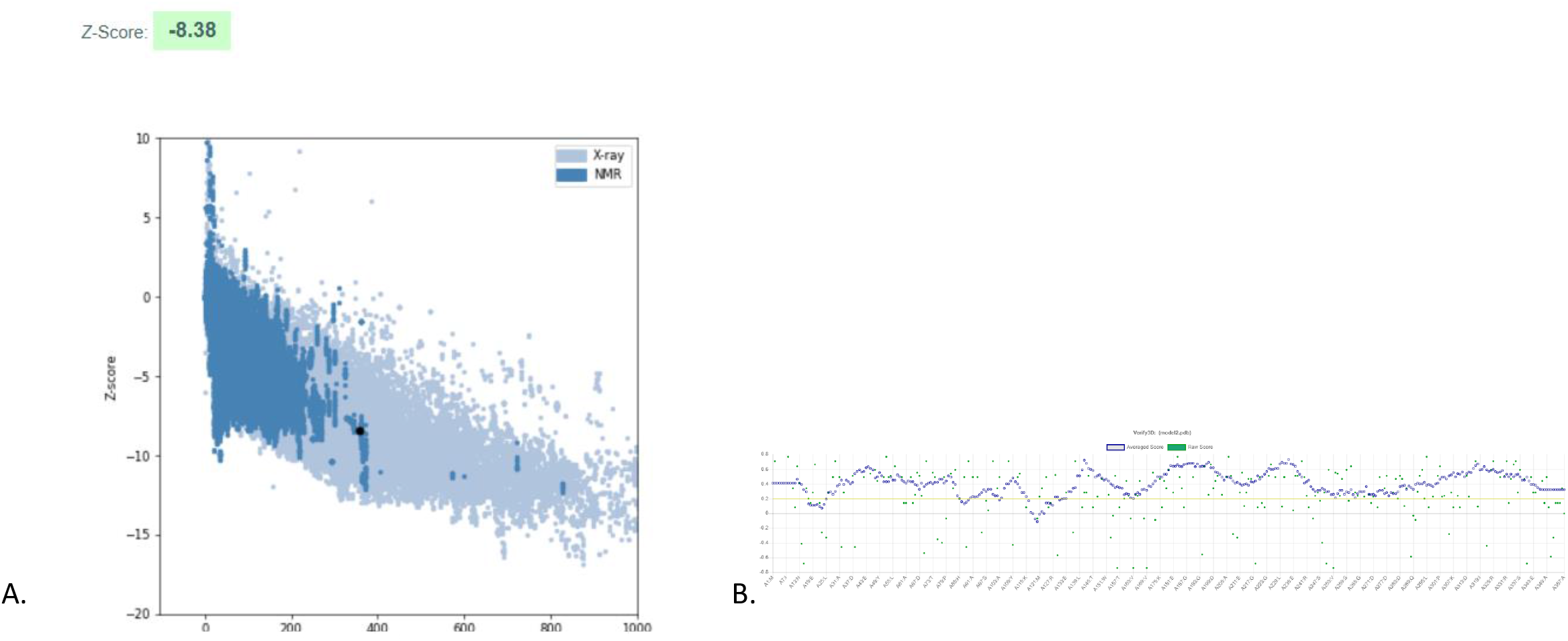
Validation of the in silico predicted model of WP_064421543.1 LLM class flavin-dependent oxidoreductase [*Mycobacterium sp*. GA-1285] by ProSa and Verify3D. A. Structure validation by ProSa, which shows the Z-score of the predicted model of Nucleoprotein (black dot), when compared to a non-redundant set of crystallographic structures (light blue dots) and NMR structures (dark blue dots). B. Structure validation by Verify3D, which shows the 3D-1D score for each atom of the predicted model of Nucleoprotein. The graphic shows that 89% of the residues of the in silico structure of Nucleoprotein presented a compatibility score of 0.2 or higher, which indicates that the structure is a high-quality model according to Verify3D.

## 4. Discussion

The study of specific kinds of mycobacterium remains to be an important endeavor for scientists interested in disease propagation due to this kind of bacterium causing some of the world’s deadliest diseases. In an effort to further this study, we studied the luciferase gene in *Mycobacterium sp. EPa45* and used for a point of reference for comparison with luciferase in other bacteria. Both the physiochemical and sequence alignment analysis showed that the *Mycobacterium sp. EPa45* was closer in relationship to pathogenic mycobacterium such as *Mycobacterium Tuberculosis* rather any other kind of non-pathogenic bacterium. *Mycobacterium sp. EPa45* has been associated with increased fatality in pulmonary disease^2^ and our discovery of it being close to pathogenic bacterium may be part of an explanation for this outcome.

A detailed distribution of the secondary structure of *Mycobacterium sp. EPa45* luciferase and their related protein gives us an idea of the spatial distribution of amino acids in the polypeptide chain. Since there was no previously available structure of *Mycobacterium sp. EPa45* luciferase, I TASSER was used to model the 3D structure of the protein. Our BlastP and CDD analysis showed that our strain of luciferase fell into the broader category of luciferase-like monooxygenases (LLMs). This kind of luciferases were found to be quite prevalent in the other bacterium studies and may be used as a benchmark for comparison.

## 5. Conclusion

Our study in the comparison of luciferase in bacterium was of a four-fold approach: (i) physiochemical characteristics, (ii) protein structure, (iii) multiple sequence alignment and (iv) phylogenetic relationships. The outcome of our study revealed that *Mycobacterium sp. EPa45* was close to pathogenic bacteria in its relationship and this gave us insight on why it caused pulmonary complications in Rhodesian patients^2^. Besides this immediate result, our study has opened a new paradigm of bacterial analysis. The similarity of Luciferase structures in pathogenic type bacterium and their differences with nonpathogenic ones indicate that there is a relationship between Luciferase structure and the risk a certain bacterium poses to patients. Given the overall association of deadly diseases with mycobacterium, our study proved to be consistent. Further study into the relationship between LLMs in different kinds of bacteria may provide a new avenue of assessment for pathogenicity.

## Reference

1. Anandan R, Dharumadurai D, Manogaran GP. An introduction to actinobacteria. In: Actinobacteria-Basics and Biotechnological Applications. Intechopen; 2016.

2. Meighen EA. Molecular biology of bacterial bioluminescence. Microbiol Mol Biol Rev. 1991;55(1):123–142.

3. Hastings JW. [13] Bacterial bioluminescence: An overview. In: Methods in Enzymology. Vol 57. Elsevier; 1978:125–135.

4. Cosma CL, Sherman DR, Ramakrishnan L. The secret lives of the pathogenic mycobacteria. Annu Rev Microbiol. 2003;57(1):641–676.

5. Pedelacq J, Nguyen MC, Terwilliger TC, Mourey L. A Comprehensive Review on Mycobacterium tuberculosis Targets and Drug Development from a Structural Perspective. Struct Biol Drug Discov Methods, Tech Pract. 2020:545–566.

6. Kato H, Ogawa N, Ohtsubo Y, et al. Complete genome sequence of a phenanthrene degrader, Mycobacterium sp. strain EPa45 (NBRC 110737), isolated from a phenanthrene-degrading consortium. Genome Announc 2015;3(4).

7. Andreu N, Zelmer A, Fletcher T, et al. Optimisation of bioluminescent reporters for use with mycobacteria. PLoS One. 2010;5(5):e10777.

8. Altschul SF, Gish W, Miller W, Myers EW, Lipman DJ. Basic local alignment search tool. J Mol Biol. 1990;215(3):403–410. doi:10.1016/S0022-2836(05)80360-2

9. Sievers F, Wilm A, Dineen D, et al. Fast, scalable generation of high-quality protein multiple sequence alignments using Clustal Omega. Mol Syst Biol. 2011;7(539). doi:10.1038/msb.2011.75

10. Kumar S, Stecher G, Li M, Knyaz C, Tamura K. MEGA X: Molecular evolutionary genetics analysis across computing platforms. Mol Biol Evol. 2018;35(6):1547–1549. doi:10.1093/molbev/msy096

11. Garg VK, Avashthi H, Tiwari A, et al. MFPPI–multi FASTA ProtParam interface. Bioinformation. 2016;12(2):74.

12. Buchan DWA, Jones DT. The PSIPRED Protein Analysis Workbench: 20 years on. Nucleic Acids Res. 2019;47(W1):W402–W407. doi:10.1093/nar/gkz297

13. Yu C, Lin C, Hwang J. Predicting subcellular localization of proteins for Gram-negative bacteria by support vector machines based on n-peptide compositions. Protein Sci. 2004;13(5):1402–1406.

14. Gasteiger E, Hoogland C, Gattiker A, Wilkins MR, Appel RD, Bairoch A. Protein identification and analysis tools on the ExPASy server. In: The Proteomics Protocols Handbook. Springer; 2005:571–607.

15. Lu S, Wang J, Chitsaz F, et al. CDD/SPARCLE: the conserved domain database in 2020. Nucleic Acids Res. 2020;48(D1):D265–D268.

16. Zhang Y. I-TASSER: Fully automated protein structure prediction in CASP8. Proteins Struct Funct Bioinforma. 2009;77(S9):100–113.

17. Yang J, Zhang Y. I-TASSER server: new development for protein structure and function predictions. Nucleic Acids Res. 2015;43(W1):W174–W181.

18. Wiederstein M, Sippl MJ. ProSA-web: Interactive web service for the recognition of errors in three-dimensional structures of proteins. Nucleic Acids Res. 2007;35(SUPPL.2):407–410. doi:10.1093/nar/gkm290

19. Laskowski RA, MacArthur MW, Moss DS, Thornton JM. PROCHECK: a program to check the stereochemical quality of protein structures. J Appl Crystallogr. 1993;26(2):283–291. doi:10.1107/s0021889892009944

20. Lovell SC, Davis IW, Adrendall WB, et al. Structure validation by C alpha geomF. Altschul, S., Gish, W., Miller, W., W. Myers, E., & J. Lipman, D. (1990). Basic Local Alignment Search Tool. Journal of Molecular Biology.etry: phi,psi and C beta deviation. Proteins-Structure Funct Genet. 2003;50(August 2002):437–450. doi:10.1002/prot.10286

